# A gradual migratory divide determines not only the direction of migration but also migration strategy of a social migrant bird

**DOI:** 10.1101/2023.02.24.529898

**Authors:** Antti Piironen, Toni Laaksonen

## Abstract

Migratory divides separate populations of migratory animals, facilitating the evolution of intraspecific differences in migration strategies. The optimal migration theory suggests differing migration strategies for birds using different flyways (with different habitats), but the knowledge regarding the impact of the flyway to the individual migration strategies is scarce. We used satellite tracking and neckband resightings to unravel the structure of the migratory divide between two European flyway populations of greylag geese *Anser anser*. We modelled satellite tracking data using Gaussian processes to study migration strategies of birds using different flyways. The mean posterior probability for an individual to migrate along the Western Flyway decreased gradually from 0.98 to 0.06 within the Baltic Sea coast, revealing a gradual migratory divide. In addition, migration strategies differed between the flyways. The birds using Western Flyway migrated earlier in autumn, performed longer annual migration and made a clear stopover during migration, whereas the birds using Central Flyway flew directly to their wintering sites. The gradual migratory divide that also divided migration strategies provides exciting insights to ecological and evolutionary factors behind migratory divides. Gaussian processes enabled modelling of detailed migration strategies, encouraging their future usage in movement ecology.

## 1. Background

Migration allows animals to use spatially and temporally versatile resources, enabling them to utilise habitats where they cannot live over the whole annual cycle (Newton, 2008). Every year billions of animals perform impressive long-distance movements tracking these periodically available resources and avoiding unfavourable environmental conditions (Thorup et al., 2017). For birds, migration often means moving between breeding sites at higher latitudes and wintering sites at lower latitudes, with some stopover sites *en route* (Newton, 2008). When the annual movements of individuals belonging to the same species are combined together, they form flyways of populations, which consist of all habitats used by the population during the annual cycle. Flyways form the basis for conservation and management of migratory species (e.g. Faaborg et al., 2010).

Flyways of birds are separated by migratory divides, contact zones on sides of which birds of the same species orientate to different directions during the non-breeding season to reach their wintering grounds (Newton, 2008). By separating flyways, migratory divides have wide-reaching evolutionary impacts. They drive intraspecific genetic differentiation and reproductive isolation (Bearhop et al., 2005; Boulet et al., 2006; Rolshausen et al., 2009) and create geographically independent, intraspecific population entities (often called management units), which form the base units for conservation and management of migratory species (Boulet et al., 2006; Faaborg et al., 2010). Importantly, migratory divides affect the availability of habitats that individuals of the same species can utilise during the non-breeding season, which can facilitate intraspecific differences in migration strategies (Alerstam & Lindström, 1990) on distinct flyways.

The structure of a migratory divide (gradual vs. precipitous) is likely to affect its ecological and evolutionary implications (for example, genetic differentiation and reproductive isolation; Delmore & Irwin, 2014). One would assume that on a continuous breeding range without any geographical obstacles, the structure of a divide would differ from a one existing alongside with a geographical barrier (such as sea, mountain or desert). However, previous studies have not tracked individuals breeding both on a divide and at different distances from it, and thus the structure of the divide has usually remained unknown (e.g. Bearhop et al., 2005; Boulet et al., 2006; Delmore et al., 2012; Hobson et al., 2015; van Bemmelen et al., 2019). This has possibly led to a simplified understanding regarding the structure of the migratory divides as only precipitous divides have been described. To better understand the ecological, evolutionary and conservation implications of migratory divides, it is vital to unravel their structures by studying migratory behaviour of individuals breeding at different distances and on a migratory divide.

While the potential importance of migratory divides on evolution of migratory behaviour has been long acknowledged, few studies have examined differences in the year-round migratory behaviours in the different flyways. The core choices for birds to be made regarding their migration are 1) the location and number of wintering and stopover sites, 2) duration of wintering period and of each stopover, and 3) timing of movement between breeding, wintering and stopover sites. These decisions can be referred to as a migration strategy (Alerstam & Lindström, 1990), which is known to have important ecological and evolutionary consequences, as well as considerable conservation and management implications (e.g. Bearhop et al., 2005; Delmore et al., 2012). Migration strategy is thought to be guided by the availability of suitable habitat (Alerstam & Lindström, 1990; Gudmundsson et al., 1991), although other factors such as weather are also known to play a role (e.g. Tøttrup et al., 2008; Tøttrup et al., 2012). According to the optimal migration theory (Alerstam & Lindström, 1990), if suitable habitats for stopovers are abundant along the flyway, one would expect to see a migration strategy consisting of frequent stopovers (refuelling periods) and short flights between the stopover sites to minimise the costs of flying with the energy stores (fat reserves). When comparing birds using distinct flyways, one would expect similar migration strategies between populations using different flyways whenever suitable habitats are equally distributed between different flyways (and different strategies if habitats are not equally distributed). Intraspecific comparisons of migration strategies between populations using different flyways (with different habitat characteristics) have produced contradicting results: Some studies have found, as expected, differing strategies (Buehler & Piersma, 2008; Delmore et al., 2012; Alves et al., 2013, van Bemmelen et al., 2019) between flyway populations. However, also surprisingly similar strategies have been found from populations using distinct flyways with different habitat characteristics (Fraser et al., 2013; Trierweiler et al., 2014), indicating that factors other than availability of suitable habitat can also contribute to migration strategies. As the conditions faced during the non-breeding season (and thus, migration strategy) are known to affect breeding populations through survival and productivity (e.g. Marra et al., 1998; Norris et al., 2004), exploring the migration strategies of populations using distinct flyways is important for a better understanding of not only the factors guiding migration strategy, but also the drivers of population dynamics of migratory populations.

Migration strategies of tracked animals are often quantified by measuring their displacement from the breeding site throughout the year (e.g. Turchin, 1998). These data are traditionally analysed by modelling them with non-linear mixed-effect models (Bunnefeld et al., 2011). These models have many obvious advantages (such as easily interpretable parameters), but they suffer from non-flexibility to model complex migration strategies. Both the intensity of individual satellite tracking and the number of animals tracked are currently increasing rapidly, allowing the exploration of new and more complex behaviours and thus, calling for novel analytical approaches (Nathan et al., 2022). Gaussian processes (GP) offer a flexible, non-parametric way to model non-linear data in the Bayesian framework, and they have been used in the machine learning community for a few decades. Although seldom used until recent years, GPs have started to gain popularity also in ecology (see e.g. Ingram et al., 2020; Wright et al., 2021; Doser et al., 2022; Piironen et al., 2022a; Wiens & Thogmartin, 2022). This is most likely due to their inherent flexibility (Rasmussen & Williams, 2006), good predictive accuracy (Ingram et al., 2020; Wright et al., 2021) and rich covariance structure that makes them an auspicious tool to model phenomena such as animal migration (Piironen et al., 2022a). Here, we demonstrate that they are a useful tool also for studying animal migration from displacement data.

The greylag geese *Anser anser* breeding in the northern Baltic Sea coast in Finland offer an excellent system to study the migratory behaviour of birds breeding in different distances from a migratory divide in a continuous landscape without geographical barriers, and to compare migratory strategies between birds breeding close to each others, but using different flyways. These birds breed on a narrow zone along the Finnish coast (Valkama et al., 2011), and they have been thought to use two different flyways during the non-breeding period: The Western Flyway (or Northwest Flyway, hereafter WF) and Central Flyway (hereafter CF, Madsen et al., 1999; Fox & Leafloor, 2018). The birds using WF breed in Fennoscandia and Western Europe, and winter sporadically in Western Europe (Nilsson, 2018). The breeding range of the birds using CF reaches from Southern Finland in the north to Czechia and Slovakia in the south, and the wintering sites are located sporadically around the Mediterranean Sea (Azafzaf et al., 2018). Despite the preliminary suggestions by Madsen et al. (1999) and Fox & Leafloor (2018), the movements of Finnish greylag geese have never been studied. Thereby, the existence of the presumed migratory divide has remained to be verified, not to mention its location. Moreover, the migration strategy of greylag geese breeding in Finland is completely unknown.

Here, we use data from satellite tracked and neckbanded individuals marked throughout the greylag goose breeding range in the Finnish Baltic Sea coast, to confirm the existence and to reveal the structure of the migratory divide. In addition to that, we analyse the daily displacements of satellite tracked individuals with Gaussian process models to compare migration strategies of birds using different flyways. In detail, we study 1) where the migratory divide is located, 2) which flyway is used by birds breeding far away, close and on the migratory divide (i.e. the structure of the divide), and 3) differences in migration strategy between birds using the different flyways. Last, we discuss the ecological, evolutionary and management implications of our results, as well as the prospects of Gaussian processes for future research on animal migration.

## 2. Material and methods

### 2.1. Field methods for satellite tracking

We caught greylag geese for satellite tracking and neckbanding throughout their breeding range in Finland in the years 2018–2022 (see Fig. I in supplemental information). The majority of the birds were caught in May and June using cannon-netting combined with short-term artificial feeding. Catching sites were located on sea shores and they were prepared prior to catching events by feeding geese with grain from several days up to some weeks. All birds marked with GPS transmitters were adults (at least two years old) and the majority of them were caught with their broods, whereas individuals from all cohorts were marked with neckbands. Additionally, we caught some birds in late June and early July, when they were flightless due to remigial moult. These birds were caught at sea with a hand-net after a short chase with a motor-driven boat. All birds were sexed using cloacal examination, aged based on the shape of wing coverts and marked also with traditional metal ring.

For satellite-tracking, we used OrniTrack-44 solar-powered GPS-GSM (Global Positioning System-Global System for Mobile communications) neckcollars produced by Ornitela UAB. The devices weigh 45 grams, which added < 2 % of the weight of the body mass of the instrumented geese. These transmitters log GPS positions and send data to the server via a GSM/GPRS network either by e-mail or SMS. To ensure the quality of the tracking data, we excluded GPS noise from the data (i.e. locations where lat 00° 00′ lon 00° 00′) and locations with hdop (horizontal dilution of precision of the GPS fix) values ≤ 2. The devices were also equipped with field-readable, three-digit code on them. Altogether, we marked 71 birds with GPS transmitters (61 females and 10 males).

We used neckbands which were made of blue, laminated and UV resistant plastic, and had white, individual three-digit field-readable codes on them. Neckbanded birds were resighted opportunistically throughout the year by voluntary observers along their flyways. We received neckband resightings from the website www.geese.org and from the database of Finnish Bird Ringing Centre.

### 2.2. Analysis of the migratory divide

To ensure the independence of observations, we excluded all birds that were known to be paired with another marked individual (e.g. birds caught as pairs as well as birds observed together with a marked individual at any point of their encounter history). As goose families move together for the first year, we also excluded all observations from birds marked as juveniles and observed during their first annual cycle. Additionally, we excluded five individuals (one satellite tracked and four neckbanded), who changed their flyway during the study period (see Discussion).

To allocate individuals into two flyways (Western or Central, hereafter WF and CF, respectively), we used the flyway range descriptions provided by Azafzaf et al. (2018) and Nilsson (2018) as following: If a neckbanded bird was resighted at least once in Denmark, northern Germany, the Netherlands, Belgium or northern France or northwest Poland, it was labelled as a WF bird. Accordingly, if a neckbanded bird was resighted east from these countries (excluding resightings from Finland), it was labelled as a CF bird. Satellite tracked birds were allocated to different flyways based on their migration routes and wintering areas following the same ranges as for neckbanded birds. We note that, based on satellite tracking data, some birds are known to use an intermediate flyway by migrating from Finnish breeding sites to stopover sites in Sweden (along the WF), but later migrating from there to the wintering sites of CF in Central Europe. Therefore, for the neckbanded birds resighted only in Sweden (*n* = 10), we considered their flyway unknown and hence excluded them from the analysis. We labelled neckbanded birds that were observed at Sweden and at the wintering sites of CF (*n* = 7) and satellite tracked with similar migratory behaviour (*n* = 2) as CF birds. For locations of satellite tracked individuals and their allocation into two flyways, see Figure 1. For the geographical distribution of all neckband resightings and their allocation into two flyways, see Figure II in supplemental information. In the end, we had 64 satellite tracked (55 females, 9 males) and 115 neckbanded (56 females, 59 males, resighted 665 times outside Finland) individuals with sufficient data for the analysis.

**Figure 1.**
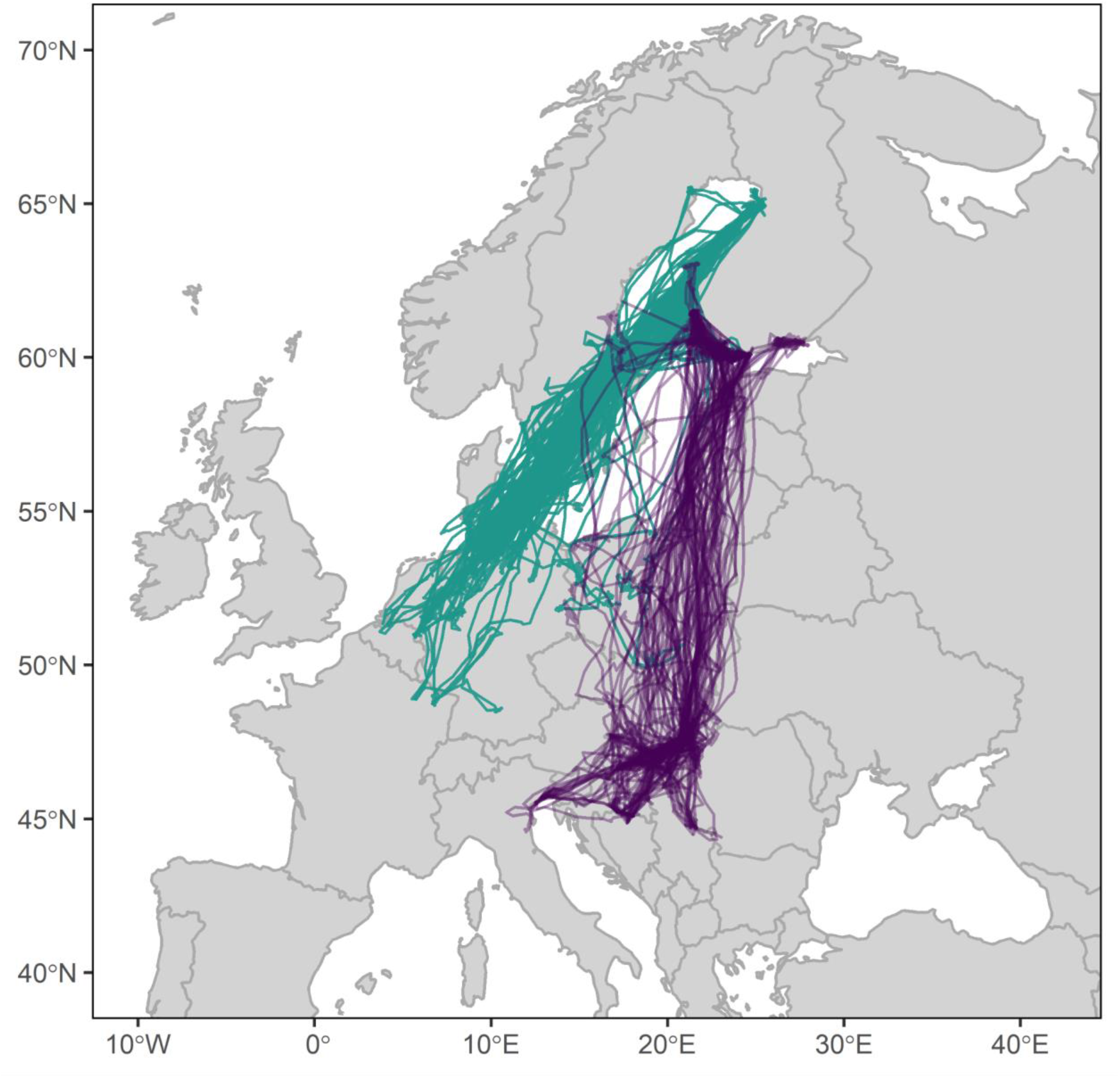
Migration routes of satellite tracked Finnish greylag geese in 2019–2022. Yearly migration routes are allocated to Western Flyway (turquoise lines) or to Central Flyway (purple lines) on their migration routes and wintering sites, following flyway range descriptions by Azafzaf et al. (2018) and Nilsson (2018). The allocation to flyways for birds that show intermediate migration routes between flyways (particularly birds that use stopover sites in Sweden (on WF) and winter in the wintering sites of CF in Central Europe) are allocated based on their wintering sites.

Following the above-mentioned allocation into two flyways, we gave the flyway status *z_i_* a value 1, if a bird was allocated to WF and a value 0, if it was allocated to the CF. We treated *z_i_* as a binomially distributed variable, i.e. *z_i_* ~ Binomial(*n*, *p*), where *p* denotes the probability for a random individual to migrate along the WF. We estimated *p* by combining binomial likelihood with uniform prior distribution *p* ~ Unif(0, 1), which is equivalent to *p* ~ Beta(1, 1). Due to conjugacy, the combination of binomial observation model and beta prior will lead to a posterior distribution *p*|*y*,*n* ~ Beta(*y*+1, *n*-*y*-1), where *y* denotes the number of individuals allocated to the WF and *n* denotes the sample size. We sampled 100,000 posterior samples for *p* in each of six coastal areas in Finland (see Results) to get the posterior distributions for *p* in each area. We analysed the data for the areas rather than as a continuous gradient to allow some clustering of marked individuals.

### 2.3. Analysis of migratory behaviour

We analysed the migratory strategies of satellite tracked birds (*n* = 64) on different flyways by measuring their daily displacement from the breeding site. We assigned each bird to either one of the flyways as described in Section 2.2. One bird changed its flyway during the study period, and hence it was assigned to different flyways in different years. We modelled the displacement data using Gaussian processes (GP), which we chose because of their flexibility (no assumptions on the form of dependence between variables are needed), because their predictive accuracy has been good in comparative studies (Ingram et al., 2020; Wright et al., 2021) and because they have appeared promising tools for modelling animal migration (Piironen et al., 2022a). For a reader new to GPs, a brief introduction in ecological context is provided in Piironen et al. (2022a), but for an in-depth introduction, see Rasmussen and Williams (2006).

In the analysis, we used satellite tracking data from the period 1.7.2019–30.6.2022 (one random location from each individual per day), and measured the displacement (*y*) for each tracked individual between the first location on 1 July in the first year the bird was tracked and every day until the tracking of the given bird ended. Due to heteroscedasticity of the data, we scaled the values *y* before the analysis so that

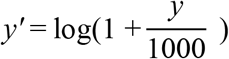

We assumed the scaled displacements *y_i_*’ to follow the gaussian observation model, i.e. *y_i_*’| *μ_i_*, *σ* ~ N(*μ_i_*, *σ*^2^). We note that, even after the scaling of *y*’, the assumption regarding the homoscedasticity of the data was not completely fulfilled. However, as the model seems to fit well to the data (see Fig. 3), we believe that this will not crucially affect the results and their interpretation. We modelled the expected value (*μ_i_*) for *y_i_*’ as a function of time (*t*) by introducing a latent function *μ*(*t*), to which we gave a zero-mean GP prior, so that *μ*(*t*) ~ *GP*(0, *k*(*t*, *t*’)). The core part of the model is the covariance function *k*(*t*, *t*’), which specifies the covariance between any *t* and *t*’. Here, we use the so-called neural network covariance function (Williams, 1998), which produces non-stationary (i.e. values of *μ*(*t*) can vary at different speeds at different values of *t*) functions and thus matches our prior expectations regarding the behaviour of *μ*(*t*). The covariance function can be written as

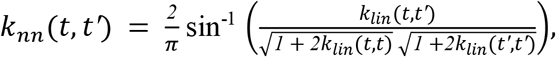

where

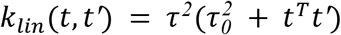

We also fitted models with quasi-periodic (see Piironen et al., 2022) and squared-exponential (see Rasmussen & Williams, 2006) covariance functions. We assessed the performance of different models using leave-one-out (LOO) cross-validation, and the model presented here performed best (see Table 1 in supplemental information for model assessment).

For model fitting, we have two hyperparameters for the covariance function (τ and τ_0_) and one hyperparameter (σ) for the likelihood to be estimated. We gave half-student-t prior distributions for τ and τ_0_ and a log-uniform prior to σ. To reduce computation time in hyperparameter estimation, we used the fully independent training and test conditional (FITC) approximation with 200 inducing points (Quiñonero-Candela & Rasmussen, 2005; Snelson & Ghahramani, 2006). We estimated hyperparameters by optimising them to their marginal maximum a posteriori values. We performed the analysis using packages adehabitatHR (displacement measurements; Calenge, 2006), gplite (fitting the GP model; Piironen, 2021) and related packages in R software version 4.1.1 (R Core Team, 2021). We included the script used in the analysis to the supplemental information.

## 3. Results

### 3.1. Migratory divide

A clear division of satellite tracked greylag geese into two flyways can be seen in Figure 1. The probabilities for individuals breeding in different coastal areas to migrate using the WF (*p*) are presented in Figure 2, in which the presence of a clear but gradual migratory divide becomes apparent. In North Ostrobothnia (at the far end of the Bothnian Bay), there is a strong statistical support that basically all birds will migrate along the WF. The probability to migrate along the WF is also high in Ostrobothnia, but decreases substantially in Satakunta (at the coast of Bothnian Sea), Uusimaa and Kymenlaakso (at the Gulf of Finland), meaning that the majority of the birds from these areas will use the CF. We note that neckband resighting data might include some misread neckbands, which can potentially bias the posterior probabilities for *p*, if the misread neckband belongs to a bird using a different flyway. However, the neckband resighting data is well in line with satellite tracking data (compare Fig. 1 and Fig. II in the supplemental information) and hence, we do not have reasons to suspect that the possible bias caused by misread neckbands would be noticeable.

**Figure 2.**
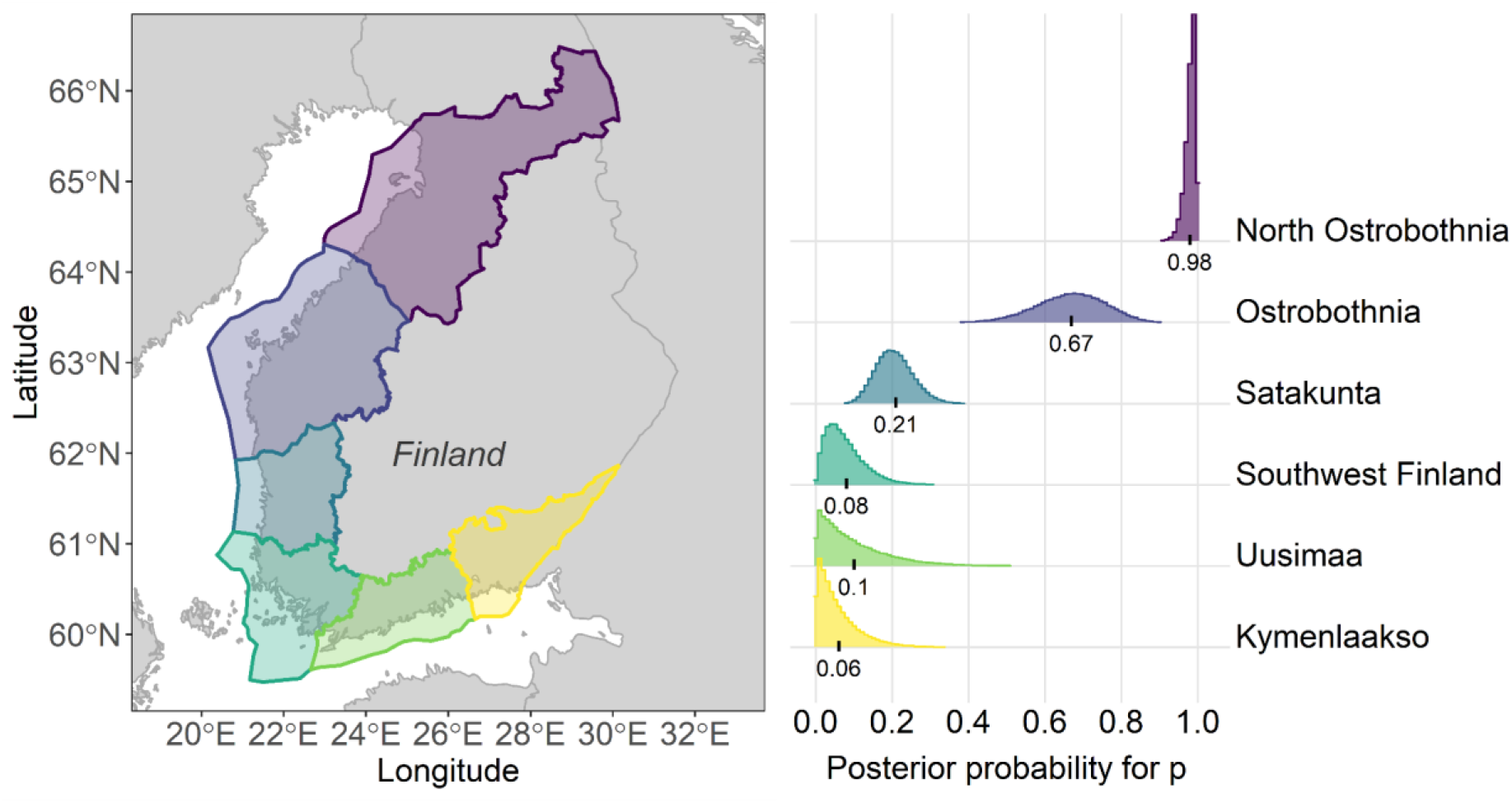
The posterior distribution for *p* (the probability for an individual to migrate along the Western Flyway) in different coastal areas in Finland. The colour of each histogram represents the probability in the area coloured with the same colour in the map. The numbers under the histograms denote the posterior mean for *p* in each county. The sample sizes for each county were *n*_North Ostrobothnia_ = 98, *n*_Ostrobothnia_ = 25, *n*_Satakunta_ = 66, *n*_Southwest Finland_ = 24, *n*_Uusimaa_ = 8 and *n*_Kymenlaakso_ = 15.

**Figure 3.**
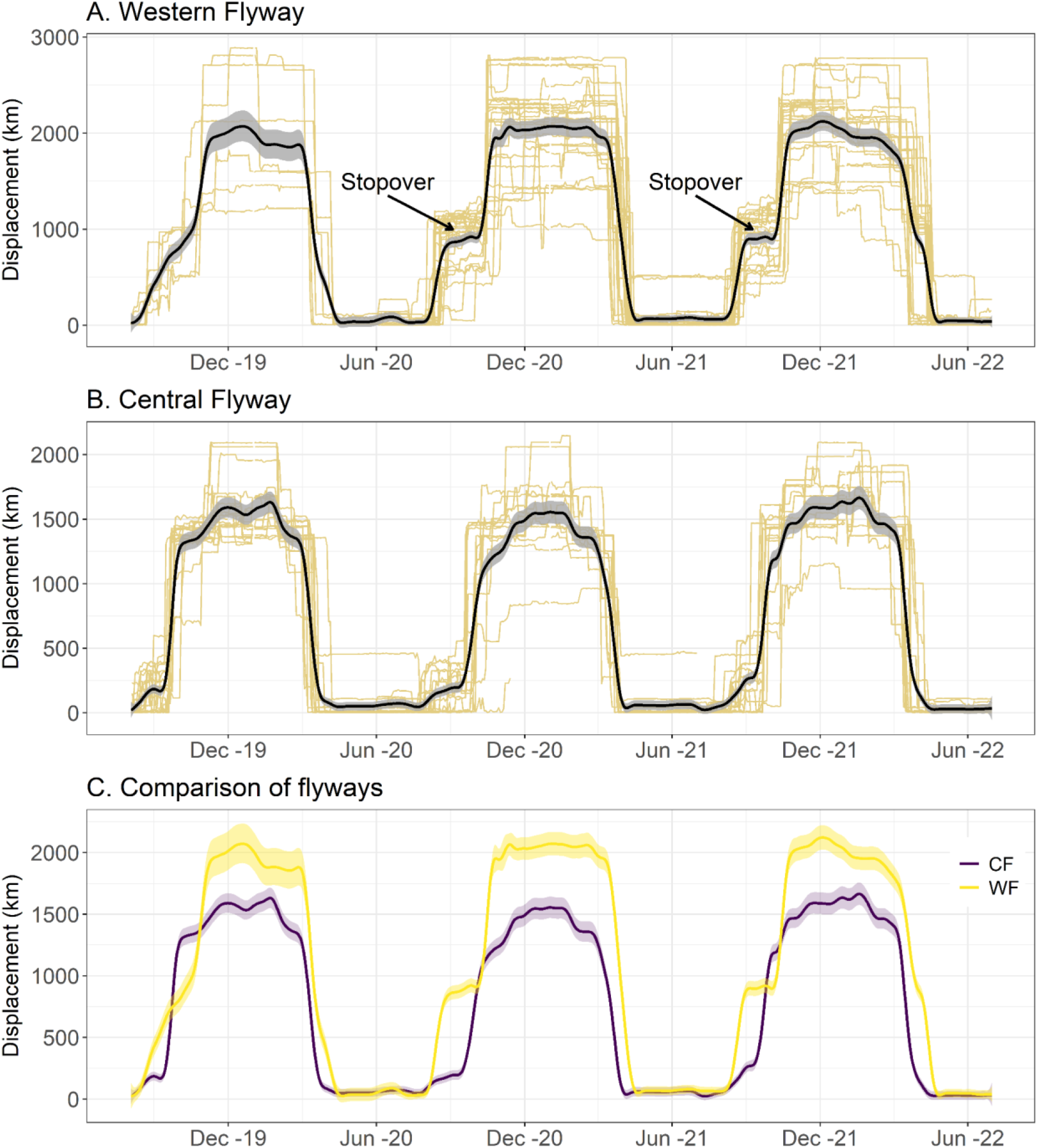
Model predictions for the displacement of satellite tracked greylag geese from their breeding sites in the Central and Western Flyways in 1.7.2019–30.6.2022. In the subplots A and B, the beige lines denote the data (i.e. displacement of each satellite tracked individual from the breeding site), and the black lines and shaded grey areas denote the posterior mean and 95 % credible interval for *μ*(*t*), respectively, all scaled to the original scale of *y* (daily displacement in kilometres from the breeding site). To facilitate easier comparison of migration strategies between flyways, subplot C visualises the above-mentioned model predictions for both flyways. Note that in all plots, the credible intervals describe the uncertainty related to the underlying function *μ*(*t*), but do not include the observation noise. Also note that the data from the Western Flyway is scarce in the year 2019, which makes the model fit also different from the subsequent years.

### 3.2. Migration strategies

Migration strategies of satellite-tracked greylag geese in the two flyways is presented in Figure 3. The overall length of the annual migration (maximum displacement between breeding and wintering grounds) is more than 500 km longer among birds using WF than among birds using CF. Birds using WF have a stopover of 1–2 months during the autumn migration, whereas birds using CF migrate relatively straight from their breeding sites to the wintering sites, disregarding some small-scale movement around breeding sites. Additionally, WF birds start their autumn migration approximately one month earlier than those using CF, whereas CF birds migrate earlier in the autumn.

## 4. Discussion

Our results show that there is a gradual migratory divide in the continuous breeding distribution of greylag geese in the Baltic Sea coast in Finland that also divides the birds to different migration strategies. The birds breeding at the far end of the Gulf of Bothnia use the Western Flyway, the birds breeding in the Gulf of Finland use the Central Flyway and the birds breeding between these two extremes scatter to the two flyways. The overall migratory journey is longer for birds using WF and they migrate earlier in the autumn (and later in the spring) than birds on CF. Birds using WF also show a clear stopover of around one month during their autumn migration, whereas CF birds migrate relatively straight from their breeding grounds to their wintering sites. These findings provide important perspectives to be considered regarding migratory divides, drivers of the migration strategy and optimal bird migration. Importantly, our study shows that GP models have features that enable modelling migratory behaviours that could not be detected with previously used modelling techniques.

Our results also provide insights for the purposes of the ongoing international management of greylag geese.

### 4.1. Evolutionary implications of the gradual migratory divide

Migratory divides are known to drive intraspecific genetic differentiation and reproductive isolation (Bearhop et al., 2005; Boulet et al., 2006; Rolshausen et al., 2009). The fact that the migratory divide among Finnish greylag geese is gradual means that a part of the birds using different flyways breed (Fig. 1) and moult (Piironen, A., unpublished data) sympatrically. This can dilute genetic differentiation between the flyways, as some genetic mixing of goose populations is known to take place during summer (probably via pair formation) on common moulting grounds (Kölzsch et al., 2019). Within the birds tracked in this study, one satellite tracked individual changed its flyway during the study period and four neckbanded birds have been resighted along both flyways, most probably indicating the change of flyway (see Fig. III and IV in supplemental information). This indicates gene flow between flyways at the overlapping breeding and moulting sites which might contribute to the low level of genetic structure among European greylag geese (Pellegrino et al., 2015). It is important to note here that pair formation among geese mainly takes place at the wintering sites (Rohwer & Anderson, 1988). The dilutive effect of a gradual migratory divide would probably be highlighted among species which form pairs mainly at the breeding sites (such as many passerines). Therefore, it is important to study the structure of the migratory divides and consider their effect on the population genetics in different species in the future.

### 4.2. Migratory behaviour of greylag geese in light of optimal migration theory

The optimal migration theory suggests that if suitable habitats for stopovers are abundant along the flyway, birds should exhibit a migration strategy consisting of frequent stopovers and short flights between the stopover sites, to minimise the costs of flying with energy stores. The migration strategies of greylag geese do not seem to follow this prediction. The greylag geese using CF fly non-stop from southern Finland to their wintering sites in Central Europe (Fig. 3). During their autumn migration, they fly over several sites in the Baltic countries and Poland that are known to be suitable stopover and wintering habitats for geese (for example, see Madsen et al., 1999; Fox & Leafloor, 2018). In addition to that, birds from both flyways migrate somewhat straight from their wintering sites to their breeding sites in spring (Fig. 3). These findings indicate that greylag geese do not try to minimise the flight with energy stores, but some other factors guide their migration strategy. Second, the birds using WF start their autumn migration approximately one month earlier than those using CF by moving from their breeding sites to stopover sites in Sweden before September (Fig. 3). Although the majority of the WF birds breed north from those using CF, the habitats and weather conditions at the Gulf of Bothnia remain suitable for geese until October-November, since other goose species (e.g. bean geese *Anser fabalis*, see Piironen et al., 2022b) occur in the area until that. Therefore, we consider it unlikely that the greylag geese would be forced to depart from the Gulf of Bothnia in August by the lack of suitable habitat as suggested by the optimal migration theory. The satellite-tracked birds indicate that hunting mortality of greylag geese is high among the birds breeding at the far end of the Gulf of Bothnia (Piironen, A., unpublished data), and thus hunting disturbance might contribute to the advanced migration schedule in the region. This, however, remains to be studied in the future. Last, the migration strategies of greylag geese differ between the WF and CF, although the habitat characteristics are at least roughly similar in both flyways. Greylag geese winter and stopover mainly in agricultural landscape holding also some wetlands (e.g. Fox & Abraham, 2017). There are more of these habitats available to birds along both flyways than used by the greylag geese (see e.g. Xu et al., 2019; d’Andrimont et al., 2021). As the optimal migration theory suggests similar migration strategies between flyways with similar habitats, our results do not indicate support for it in this sense. Although we have not quantified the availability and quality of habitat in each flyway or explained the differences in migration strategies with quantitative habitat factors, we consider that differences in habitat characteristics won’t probably explain the observed differences in migration strategies between the flyways. To better understand bird migration and how migratory birds can respond to habitat loss and environmental changes, factors guiding migration strategies should be unravelled in future studies.

### 4.3. Gaussian processes in modelling animal movement using displacement data

We modelled migratory behaviour using Gaussian processes (GP) instead of non-linear mixed-effect model (Bunnefeld et al., 2011), which has been the most common choice for this kind of analysis. The flexibility of GPs appeared beneficial in finding fine-scale migratory behaviour such as stopover during migration, which would have been impossible to model with commonly used methods (i.e. non-linear mixed-effect models) as their fit is a double sigmoid curve. In addition to that, the possibility to implement periodic (or quasi-periodic, see Piironen et al., 2022a) covariance structure to the model is many times beneficial when modelling phenomena such as animal migration over multiple years (as migration patterns in different years often remind each other, but are not exactly similar). However, the drawback of GPs in this context is the interpretation of the parameters as their hyperparameters are very difficult to interpret in an ecologically meaningful way (as opposed to the model presented by Bunnefeld et al., (2011), which provides easily interpretable parameters). However, GPs have proven to be a promising tool for multiple non-linear problems in ecology, and their capabilities should be better explored and utilised in the expanding field of movement ecology. As satellite tracking will most likely continue to increase its popularity among movement ecologists in the future, it would be useful to conduct comparative studies of different techniques in modelling displacement data to unravel the best methods to analyse animal migration.

### 4.4. Implications for population delineation and international management

Finland is a range state in the international management of the Northwest-Southwest European population of greylag geese (which uses the WF), but the proportion of Finnish greylag geese belonging to this population has been unknown (Bacon et al., 2019). Our results enable the allocation of Finnish breeding population between the flyway populations based on their breeding grounds (Fig. 2). Additionally, Finnish greylag geese have formerly been observed to winter as far south as in Spain (along WF) or in North Africa (along CF, Andersson et al., 2001). Our data therefore indicates that wintering sites have shifted northwards since that (Fig. 1). Although the exploration of spatio-temporal distribution of the population is beyond the scope of this study, our results indicate similar shifts among Finnish greylag geese than the ones that have recently been observed among greylag geese breeding in Sweden (Månsson et al., 2022), and are most likely mainly caused by climate change. As the climate will probably continue warming also in the future and greylag geese have shown their ability to rapidly adapt to the changing environmental conditions, it will remain important to track the changing migration patterns also in the future.

## Ethics statement

Capturing and marking of birds was done by the approval of Finnish Wildlife Agency (licence number 2019-5-600-01158-8).

## Conflict of interest statement

None declared.

## Funding statement

This work was funded by the Finnish Wildlife Agency by a grant to authors. Additionally, Finnish Wildlife Agency, Natural Resource Institute Finland (Luke), Finnish Hunter’s Association, Uittokalusto Oy, Swarovski Optik, Kartanon Riista Oy, G2 Invest Oy, Raseborg Stad and relatives of Jaana Ylönen funded satellite tracking devices. Field work was funded by Luonnon-ja Riistanhoitosäätiö and Suomen Riistanhoito-Säätiö by grants to AP.

## Author’s contributions

AP conceived the original idea for the study, acquired funding, designed the study, led the field work, analysed the data and led the manuscript writing. TL supervised structural decisions and participated in the manuscript writing. Both authors agreed to the final version of the manuscript.

## Acknowledgements

We thank Juho Piironen for statistical advice and Tuomas Seimola for providing Luke’s satellite tracking data. We thank Noora Metsäranta, Juho Lukka, Tatu Hokkanen, Jorma Nurmi, Harri Forsten, Aki S. Perälä, Eero Rautanen (+family), Allan and Petri Siironen, Niko Jyrkkä, Jani Luosujärvi, Antti Toppila, Auvo Arffman, Mika and Tapio Majava, Peter Wollsten, Jörgen and Axel Hermansson, Ari Isosalo, Jouni Kannonlahti, Juha Pikkarainen, Mikko Haapoja, Henri Selin, Kaj Sjögård, Mika Karvonen, Heikki Vilponen, Sari Holopainen, Samuli Karppinen, Henri Kopisto, Aku Törmikoski, Ismo Lehkonen and Jarno Ruohonen for help in the field work. We also thank Finnish Bird Ringing Centre and the administrators of www.geese.org for their contributions with the neckband resighting data. Last, we thank all birdwatchers and hunters who submitted their neckband resightings.

## Notes

### Competing Interest Statement

The authors have declared no competing interest.

## References

Alerstam T, Lindström Å. 1990 Optimal bird migration: The relative importance of time, energy and safety. In Bird migration: The physiology and ecophysiology (ed. Gwinner E), Berlin, Germany: Springer.

Alves JA et al. 2013 Costs, benefits, and fitness consequences of different migratory strategies. Ecology. 94, 11–17. (doi:10.1890/12-0737.1)

Andersson Å, Follestad A, Nilsson L, Persson H. 2001 Migration patterns of Nordic greylag geese *Anser anser*. Ornis Svec. 11, 19–58. (doi:10.34080/os.v11.22859)

Azafzaf H, Baccetti N, Smart M, Musil P, Rozenfeld S, Samraoul B. 2018 E4 Central Europe/North Africa greylag goose *Anser anser*. In A global audit of the status and trends of Arctic and northern hemisphere goose populations (component 2: population accounts) (ed. Fox AD, Leafloor JO, Akureyri, Iceland: Conservation of Arctic Flora and Fauna International Secretariat.

Bacon L et al. 2019 Spatio-temporal distribution of greylag goose *Anser anser* resightings on the north-west/south-west European flyway: guidance for the delineation of transboundary management units. Wildlife Biol. wlb.00533. (doi:10.2981/wlb.00533)

Bearhop S et al. 2005 Assortative mating as a mechanism for rapid evolution of a migratory divide. Science. 310, 502–504. (doi:10.1126/science.1115661)

Boulet M, Gibbs HL, Hobson KA. 2006 Integrated analysis of genetic, stable isotope, and banding data reveal migratory connectivity and flyways in the northern yellow warbler *(Dendroica petechia;* Aestiva group). Ornithol. Monogr. 61, 29–78. (doi:10.2307/40166837)

Buehler DM, Piersma T. 2008 Travelling on a budget: Predictions and ecological evidence for bottlenecks in the annual cycle of long-distance migrants. Proc. Royal Soc. B. 363, 247–266. (doi:10.1098/rstb.2007.2138)

Bunnefeld N et al. 2011 A model-driven approach to quantify migration patterns: Individual, regional and yearly differences. J. Anim. Ecol. 80, 466–476. (doi:10.1111/j.1365-2656.2010.01776.x)

Calenge C. 2006 The package “adehabitat” for the R software: A tool for the analysis of space and habitat use by animals. Ecol. Modell. 197, 516–519. (doi: 10.1016/j.ecolmodel.2006.03.017)

d’Andrimont R, Verhegghen A, Lemoine G, Kempeneers P, Meroni M, van der Velde M. 2021 From parcel to continental scale – A first European crop type map based on Sentinel-1 and LUCAS Copernicus in-situ observations. Remote Sens. Env. 266, 112708. (doi:10.1016/j.rse.2021.112708)

Delmore KE, Irwin DE. 2014 Hybrid songbirds employ intermediate routes in a migratory divide. Ecol. Lett. 17, 1211–1218. (doi:10.1111/ele.12326)

Delmore KE, Fox JW, Irwin DE. 2012 Dramatic intraspecific differences in migratory routes, stopover sites and wintering areas, revealed using light-level geolocators. Proc. Royal Soc. B. 279, 4582–4589. (doi:10.1098/rspb.2012.1229)

Doser JW, Finley AO, Kery M, Zipkin EF. 2022 spOccupancy: An R package for single-species, multi-species, and integrated spatial occupancy models. Methods Ecol. Evol. 13, 1670–1678. (doi:10.1111/2041-210X.13897)

Faaborg J et al. 2010 Conserving migratory land birds in the new world: Do we know enough? Ecol. Appl. 20, 398–418. (doi:10.1890/09-0397.1)

Fox AD, Abraham KF. 2017 Why geese benefit from the transition from natural vegetation to agriculture. Ambio. 46 (Suppl. 2), S188–S197. (doi:10.1007/s13280-016-0879-1)

Fox AD, Leafloor JO. 2018 A global audit of the status and trends of Arctic and Northern Hemisphere goose populations (component 2: population accounts). Akureyri, Iceland: Conservation of Arctic Flora and Fauna International Secretariat.

Fraser KC et al. 2013 Consistent range-wide pattern in fall migration strategy of purple martin *(Progne subis),* despite different migration routes at the Gulf of Mexico. Auk. 130, 291–296. (doi:10.1525/auk.2013.12225)

Gudmundsson GA, Lindström Å, Alerstam T. 1991 Optimal fat loads and long-distance flights by migrating knots *Calidris canutus*, sanderlings *C. alba* and turnstones *Arenaria interpres*. Ibis. 133, 140–152. (doi:10.1111/j.1474-919X.1991.tb04825.x)

Hobson KA et al. 2015 A continent-wide migratory divide in North American breeding barn swallows *(Hirundo rustica)*. PLoS ONE. 10, e0129340. (doi:10.1371/journal.pone.0129340)

Ingram M, Vukcevic D, Golding N. 2020 Multi-output Gaussian processes for species distribution modelling. Methods Ecol. Evol. 11, 1587–1598. (doi:10.1111/2041-210X.13496)

Kölzsch A et al. 2019 Flyway connectivity and exchange primarily driven by moult migration in geese. Mov. Ecol. 7, 1–11. (doi:10.1186/s40462-019-0148-6)

Madsen J, Fox AD, Cracknell G. 1999 Goose populations of the Western Palearctic. A review of status and distribution, 1st edn. Wageningen, the Netherlands: Wetlands International.

Marra PP, Hobson KA, Holmes RT. 1998 Linking winter and summer events in a migratory bird by using stable-carbon isotopes. Science. 282, 1884–1886. (doi:10.1126/science.282.5395.1884)

Månsson J, Liljebäck N, Nilsson L, Olsson C, Kruckenberg H, Elmberg J. 2022 Migration patterns of Swedish greylag geese *Anser anser* – implications for flyway management in a changing world. Eur. J. Wild. Res. 68, 15. (doi:10.1007/s10344-022-01561-2)

Nathan R et al. 2022 Big-data approaches lead to an increased understanding of the ecology of animal movement. Science. 375, 734. (doi:10.1126/science.abg1780)

Newton I. 2008 The migration ecology of birds. London, UK: Elsevier https://doi.org/10.1016/b978-0-12-517367-4.x5000-1.

Nilsson L. 2018 E3 Northwest European Greylag Goose *Anser anser*. In A global audit of the status and trends of Arctic and northern hemisphere goose populations (component 2: population accounts) (ed. Fox AD, Leafloor JO, Akureyri, Iceland: Conservation of Arctic Flora and Fauna International Secretariat.

Norris DR, Marra PP, Montgomerie R, Kyser TK, Ratcliffe LM. 2004 Reproductive effort, molting latitude, and feather color in a migratory songbird. Science. 306, 2249–2250. (doi:10.1126/science.1103542)

Pellegrino I, Cucco M, Follestad A, Boos M. 2015 Lack of genetic structure in greylag goose (*Anser anser*) populations along the European Atlantic flyway. PeerJ. 3, e1161. (doi:10.7717/peerj.1161)

Piironen A, Piironen J, Laaksonen T. 2022 Predicting spatio-temporal distributions of migratory populations using Gaussian process modelling. J. Appl. Ecol. 59, 1146–1156. (doi:10.1111/1365-2664.14127)

Piironen A, Fox AD, Kampe-Persson H, Skyllberg U, Therkildsen OR, Laaksonen T. 2022 When and where to count? Implications of migratory connectivity and nonbreeding distribution to population censuses in a migratory bird population. Popul. Ecol. early access. (doi:10.1002/1438-390X.12143)

Piironen J. 2021 gplite: General purpose Gaussian process modelling. R package.

Quiñonero-Candela J, Rasmussen CE. 2005 A unifying view of sparse approximate Gaussian process regression. J. Mach. Learn. Res. 6, 1939–1959.

R Core Team. 2021 R: A language and environment for statistical computing. Vienna, Austria: R Foundation for Statistical Computing.

Rasmussen CE, Williams CKI. 2006 Gaussian processes for machine learning. Massachusetts, United States: MIT Press.

Rohwer FC, Anderson MG. 1988 Female-biased philopatry, monogamy, and the timing of pair formation in migratory waterfowl. In Current Ornithology 5 (ed. Johnston RF), Boston, United States: Springer.

Rolshausen G, Segelbacher G, Hobson KA, Schaefer HM. 2009 Contemporary evolution of reproductive isolation and phenotypic divergence in sympatry along a migratory divide. Curr. Biol. 19, 2097–2101. (doi:10.1016/j.cub.2009.10.061)

Snelson E, Ghahramani Z. 2006 Sparse Gaussian processes using pseudo-inputs. In Advances in neural information processing systems (ed. Weiss Y, Schölkopf B, Platt JC, Massachusetts, US: MIT Press.

Thorup K et al. 2017 Resource tracking within and across continents in long-distance bird migrants. Sci. Adv. 3, 1–11. (doi:10.1126/sciadv.1601360)

Tøttrup AP et al. 2012 Drought in Africa caused delayed arrival of European songbirds. Science. 338, 1307. (doi:10.1126/science.1227548)

Tøttrup AP, Thorup K, Rainio K, Yosef R, Lehikoinen E, Rahbek C. 2008 Avian migrants adjust migration in response to environmental conditions *en route*. Biol. Lett. 4, 685–688. (doi:10.1098/rsbl.2008.0290)

Trierweiler C et al. 2014 Migratory connectivity and population-specific migration routes in a long-distance migratory bird. Proc. Royal Soc. B. 281, 20132897. (doi:10.1098/rspb.2013.2897)

Turchin P. 1998 Quantitative Analysis of Movement. Sunderland, UK: Sinauer Associates.

Valkama J, Vepsäläinen V, Lehikoinen A. 2011 Finnish breeding bird atlas III. Helsinki, Finland: Finnish Museum of Natural and the Ministry of Environment.

van Bemmelen RSA et al. 2019 A migratory divide among red-necked phalaropes in the Western Palearctic reveals contrasting migration and wintering movement strategies. Front. Ecol. Evol. 7, 86. (doi:10.3389/fevo.2019.00086)

Wiens AM, Thogmartin WE. 2022 Gaussian process forecasts *Pseudogymnoascus destructans* will cover coterminous United States by 2030. Ecol. Evol. 12, e9547. (doi:10.1002/ece3.9547)

Williams CKI. 1998 Computation with infinite neural networks. Neural Comput. 10, 1203–1216.

Wright WJ, Irvine KM, Rodhouse TJ, Litt AR. 2021 Spatial Gaussian processes improve multispecies occupancy models when range boundaries are uncertain and nonoverlapping. Ecol. Evol. 11, 8516–8527. (doi:10.1002/ece3.7629)

Xu T, Weng B, Yan D, Wang K, Li X, Bi W, Li M, Cheng X, Liu Y. 2019 wetlands of international importance: status, threats, and future protection. Int. J. Environ. Res. Public Health. 16, 1818. (doi:10.3390/ijerph16101818)

