## Supplement 1 for "A gradual migratory divide determines not only the direction of migration but also migration strategy of a social migrant bird"

### SUPPLEMENTAL INFORMATION

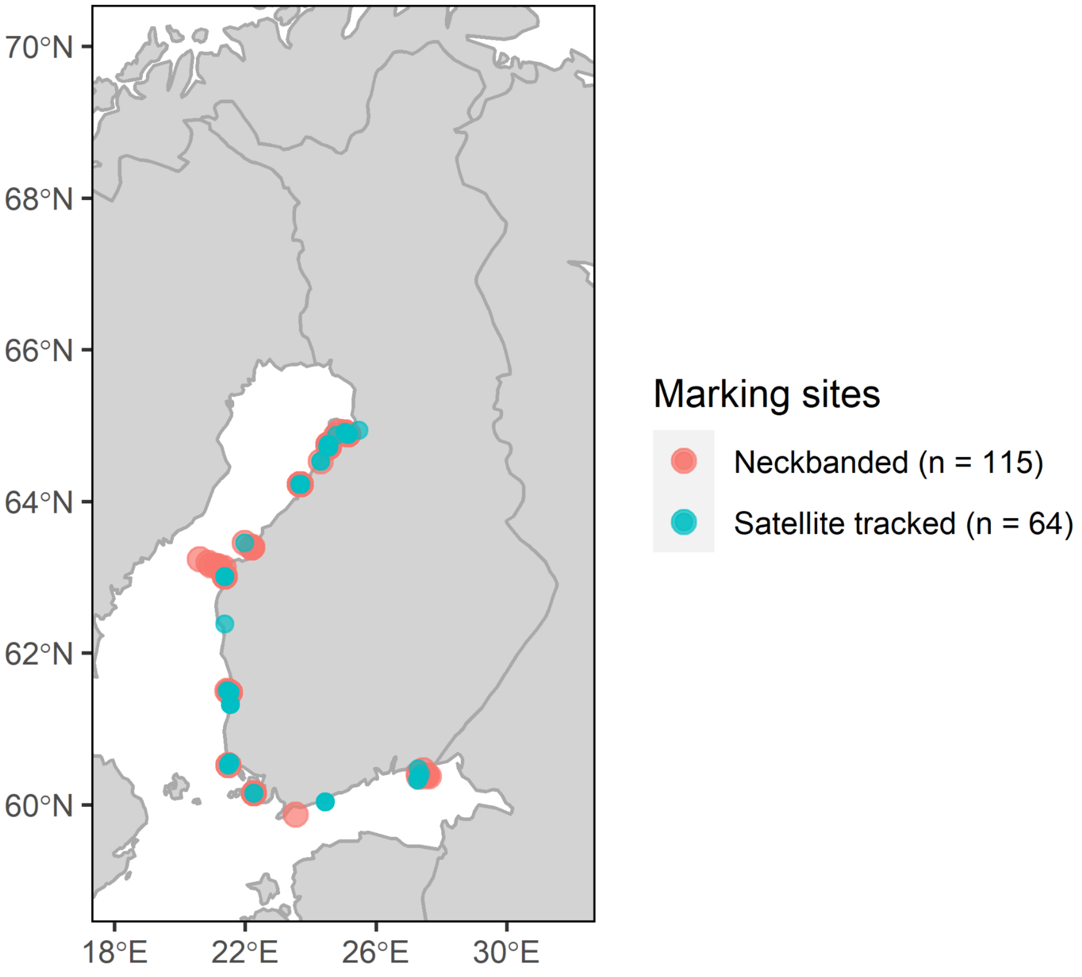


Figure I. Marking sites for greylag geese marked in Finland 2018–2022 and used in this study.

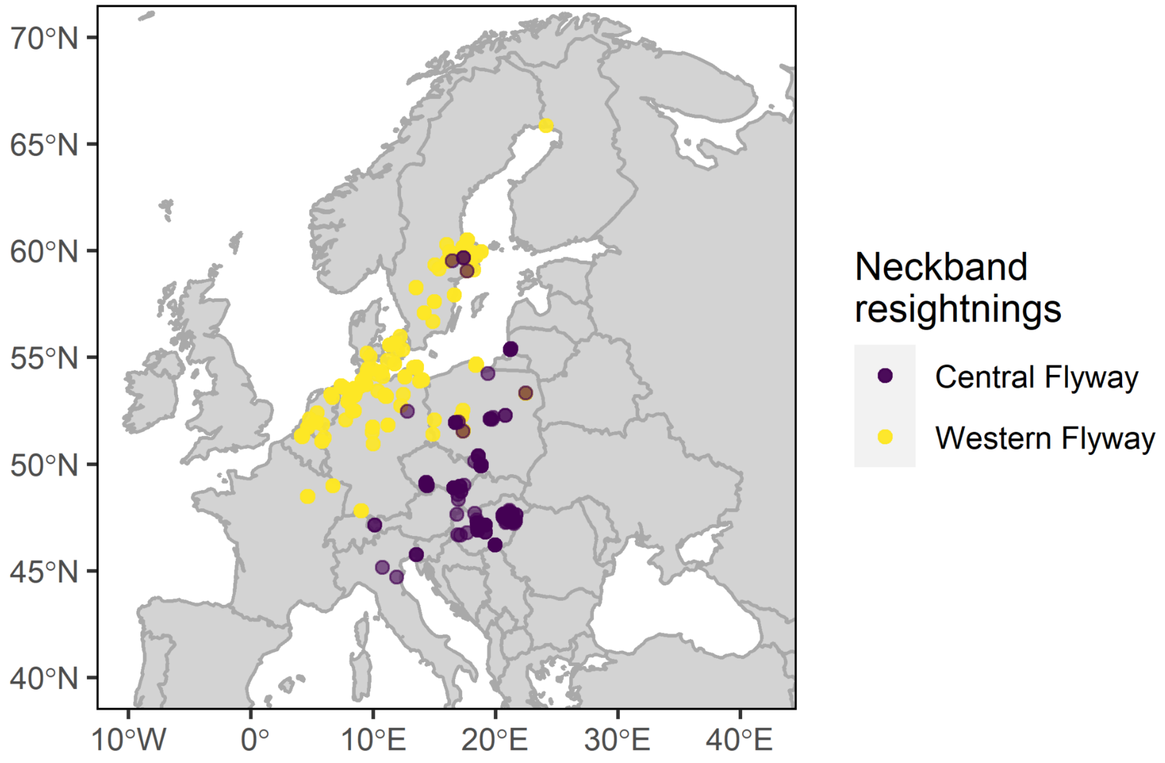


Figure II. Distribution of neckband resightings outside Finland for birds allocated to WF (yellow dots) and CF (green dots) following flyway range descriptions provided by Azafzaf et al. (2018), and Nilsson (2018). Purple dots in Sweden represent birds that have been observed on stopover sites in Sweden (on WF) and in the wintering grounds in Central Europe (on CF, see also Figure 1 in the article).

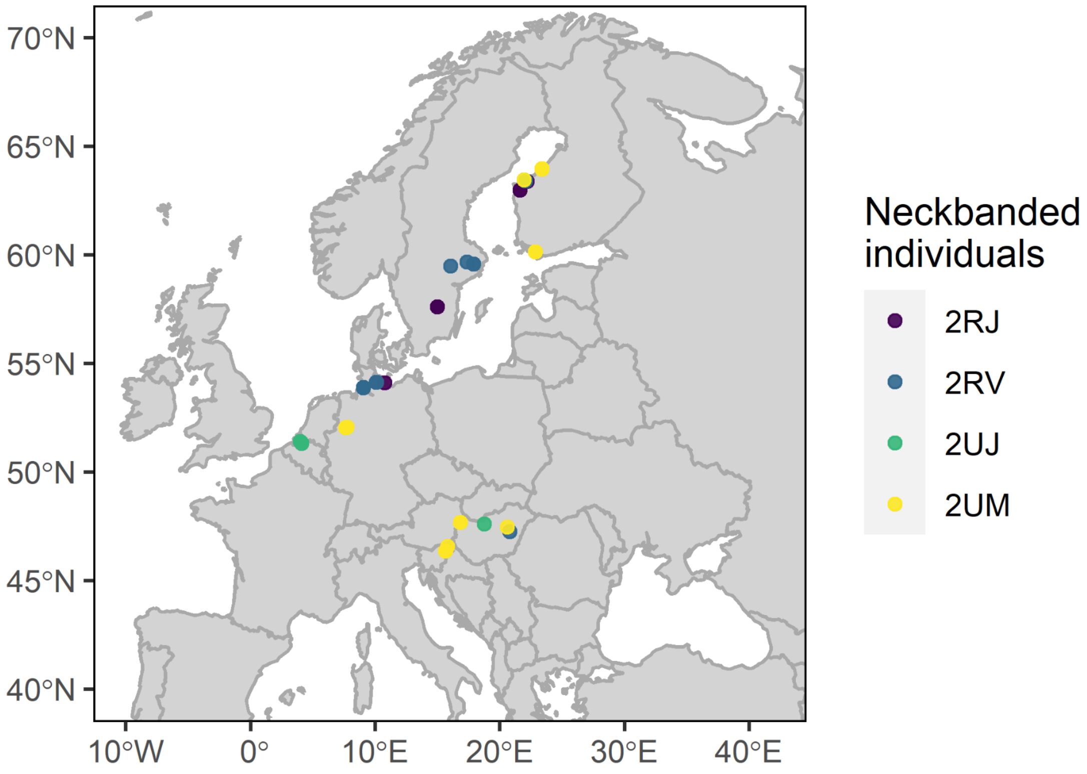


Figure III. Neckband resightings from birds which were resighted on both flyways during the study period 2018–2022.

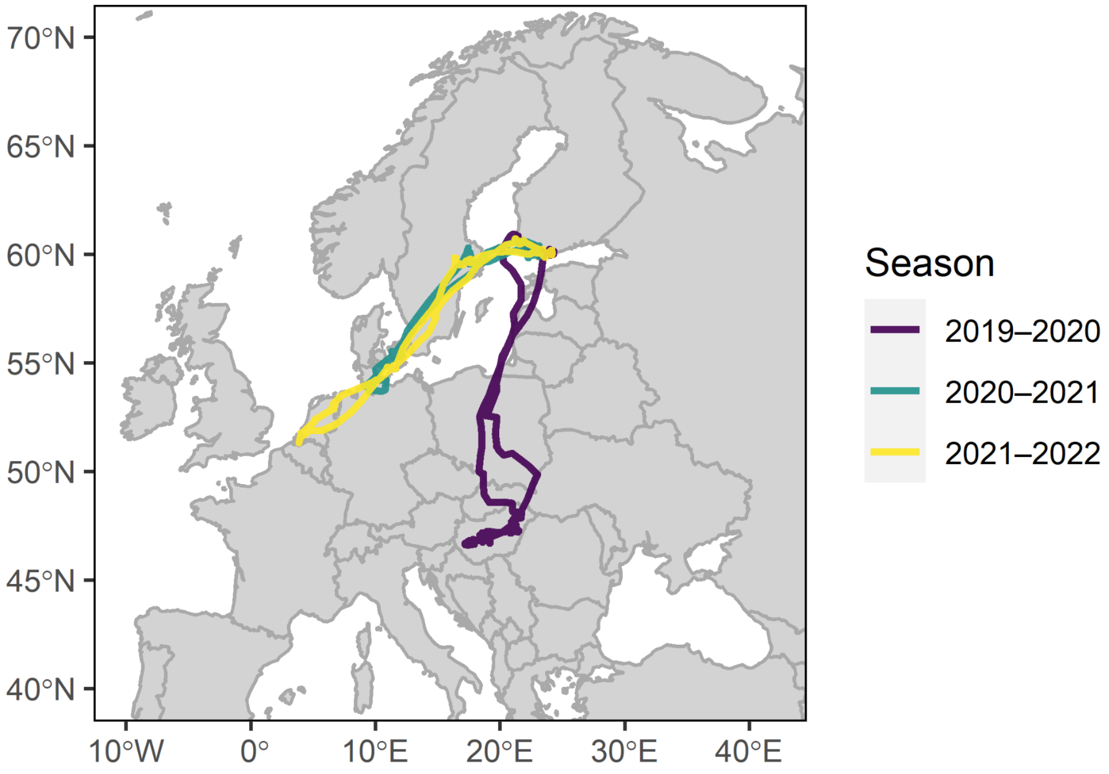


Figure IV. Seasonal tracks of the satellite tracked individual which changed its flyway during the study period in summer 2020. Each track with a unique colour represents a track between 1 July and 30 June in given years, i.e. one migration from the breeding sites to wintering sites and back.

Table 1. Model assessment. First column shows the different models i.e. the models with different covariance functions (kernels), the second column shows the difference in the sum of leave-one-out (LOO) cross-validation log-predictive densities (with the standard error in the third column) for pairwise comparison to chosen (best) model. Positive values indicate better and negative values worse performance than the chosen model.

| Model | LOO difference | SD |
|---|---|---|
| *Western Flyway* | | |
| Model with squared-exponential kernel | -21.555589 | 7.075035 |
| Model with quasi-periodic kernel | -3.288889 | 2.976757 |
| Model with neural network kernel | 0.000000 | 0.000000 |
| *Central Flyway* | | |
| Model with squared-exponential kernel | -37.358402 | 8.616063 |
| Model with quasi-periodic kernel | -9.210113 | 4.547915 |
| Model with neural network kernel | 0.000000 | 0.000000 |
